# Phylogeny and Metadata Network Database for Epidemiologic Surveillance

**DOI:** 10.1101/2022.04.19.488067

**Authors:** Garrick Stott, Leke Lyu, Gabriella Veytsel, Jacky Kuo, Ryan Lewis, Armand Brown, Kayo Fujimoto, Justin Bahl

**Affiliations:** Center for Ecology of Infectious Diseases, Institute of Bioinformatics, University of Georgia Athens, GA 30602, USA; Department of Health Promotion Behavioral Sciences, School of Public Health, The University of Texas Health Science Center at Houston, Houston, TX 77030, USA; Bureau of Epidemiology, Houston Health Department, Houston, TX 77054, USA; Department of Infectious Diseases, Department of Epidemiology and Biostatistics, University of Georgia, Athens, GA 30602, USA

**Keywords:** SARS-CoV-2, Database Management Systems, Phylogenetic Relationships, Molecular Epidemiology

## Abstract

The ongoing SARS-CoV-2 pandemic has highlighted the difficulty in integrating disparate data sources for epidemiologic surveillance. To address this challenge, we have created a graph database to integrate phylogenetic trees, associated metadata, and community surveillance data for phylodynamic inference. As an example use case, we divided 22,713 SARS-CoV-2 samples into 5 groups, generated maximum likelihood trees, and inferred a potential transmission network from a forest of minimum spanning trees built on patristic distances between samples. We then used Cytoscape to visualize the resultant graphs.

## 1 Introduction

Integration of phylogenetic, epidemiologic, and contact tracing networks provides insights into transmission dynamics and aids in cluster identification, but this approach is still a nascent field. Many tools available to integrate these data for a single analysis such as MicrobeTrace and StrainHub focus on providing a visualization tool for end users or are limited by local hardware.[1, 2] COVID-19 presents a unique challenge due to the sheer volume of both sequencing data and metadata. There is a need for an infrastructural approach to integrate these data to ensure they are scalable in long-term surveillance efforts. We propose a graph database as a source of record to integrate these data and identify transmission clusters at scale. Graph databases, a type of NoSQL databases, differ from relational database management systems (RDBMS) in that data are stored as nodes, relationships, and properties rather than a series of tables joined together. The primary benefit from such a design is that computation involving primarily relationships between entities are easily traversed, making network computations more efficient through indexes along these paths.[3]

## 2 Methods

### 2.1 Phylogeny Storage

Trees are stored in the graph database as a collection of tree alignment graphs (TAG), an approach developed as part of the Open Tree of Life Project, allowing us to share metadata across multiple trees with potentially conflicting relationships. A TAG is a collection of nodes, representing least inclusive common ancestors (LICA) and samples, joined by a series of edges indicating child relationships indexed by source id.[4] Figure 1 provides an example pair of trees and how they would be stored in the graph database. Each Newick file is parsed by a python script which generates a CSV file of edges in the TAG. A cypher script then reads the CSV file into the graph database, linking leaf nodes to their associated metadata and aligning each tree. Storing trees in this way will allow us to extract the original tree, subsets of each tree, or supersets of trees as needed. Metadata are stored either as nodes or properties within a node depending on whether they form a community of interest (e.g. a network of location data and related samples) or used to modify node attributes in a visualization (e.g. coloring nodes by clade or race).

**Figure 1:**
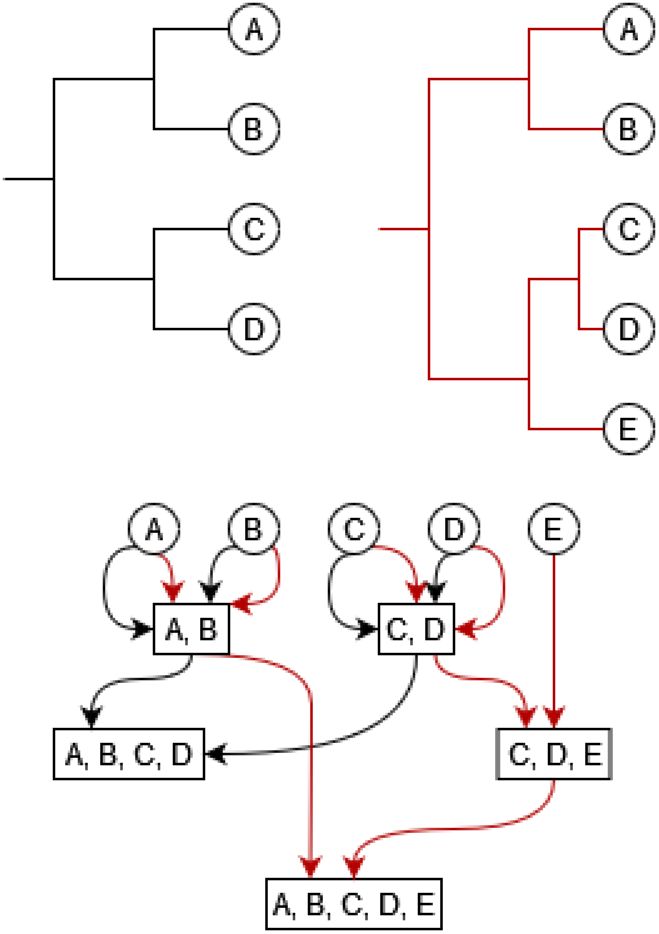
Example mapping schematic. Two source trees with partially overlapping taxon sets (top) are mapped to a graph (bottom). The colored edges of the graph correspond to the source trees. Internal graph nodes, representing least inclusive common ancestors (LICA), are represented as rectangles and terminal taxa are represented as circles.

### 2.2 Transmission Network Inference

We calculated the patristic distance between each sample for each tree, which provided a simulation of periodic updates to a phylogenetic tree as samples are collected. While pairwise genetic distance is often used in transmission clustering due to its low computational cost, patristic distances use more of the information available allowing for easier differentiation between rapidly and slowly evolving sites.[5] From the patristic distance networks, we then generated a forest of minimum spanning trees (MST), each starting from a different initial node collected August 1, 2021. While distance thresholds can be used to infer transmission, MSTs are less prone to discarding links between samples. We then aggregated the resultant edges (as counts and average patristic distances) to build our inferred transmission network. While molecular sequence data alone is insufficient to truly infer transmission events given the rapid transmission of SARS-CoV-2, we plan to leverage these networks to connect disjoint contact tracing and venue affiliation data. Figure 2 shows the final graph schema.

**Figure 2:**
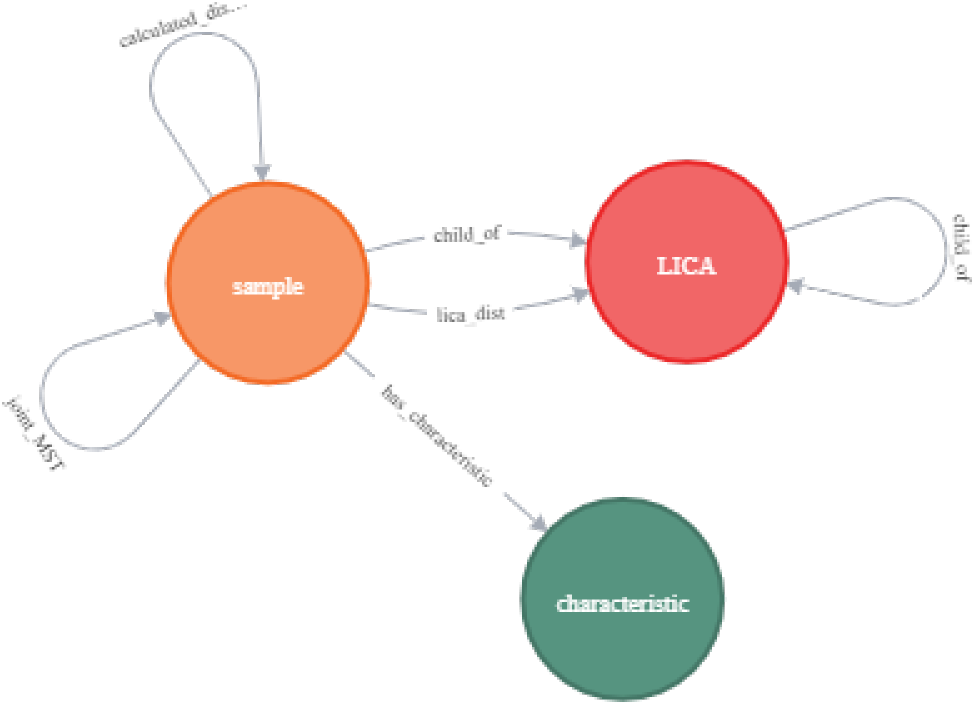
Final graph schema. Sample and LICA nodes both also share a phylogeny label to aid in indexing.

### 2.3 Visualization Tools

Once the database has been loaded and analytical queries have been executed, there are several options for visualizing results. Users can install Neo4j Bloom on the graph database for simple exploration of the data. This tool is the easiest to set up, but we found that it rapidly reaches a ceiling after just 125,000 nodes and relationships. Another option is to use Gephi, but to use this tool, users need to start listening for data streams in Gephi before executing an APOC graph streaming query to push the data to the app.[6] Alternatively, users can connect using the Neo4j plugin for Cytoscape.[7] We found this option to be the most intuitive and sustainable for ad-hoc visualization since you can remotely connect to the graph using a read only user account on the database. Finally, developers can connect directly to the REST API in their programming language of choice.

## 3 Example use case

We obtained metadata from 22,713 samples collected August-September 2021 in Texas and downloaded full sequences from GISAID for each sample. We aligned sequences to the Wuhan-Hu-1 reference genome using MAFFT 7.313, then split the dataset into 5 subsets to mimic an ongoing pandemic response with biweekly updates: 08/01/2021-08/14/2021, 08/01/2021-08/28/2021, 08/01/2021-09/11/2021, 08/01/2021-09/25/2021, 08/01/2021-09/30/2021. For each subset, we generated a maximum likelihood (ML) tree under a generalized time-reversible model of nucleotide evolution using FastTree 2.1.11. These five trees were then loaded into our graph database as TAGs and connected to their associated metadata before generating a network of MSTs for each tree.

For each of the 5 data subsets, we generated a graph of samples connected by their aggregated MST network with edge weights determined by percentage of MSTs that share the edge. We then used Cytoscape to connect to and visualize the network of minimum spanning trees generated in our analysis, seen in Figure 3. These data were then also used to generate a geospatial transmission network from sample zip codes, shown in figure 4.

**Figure 3:**
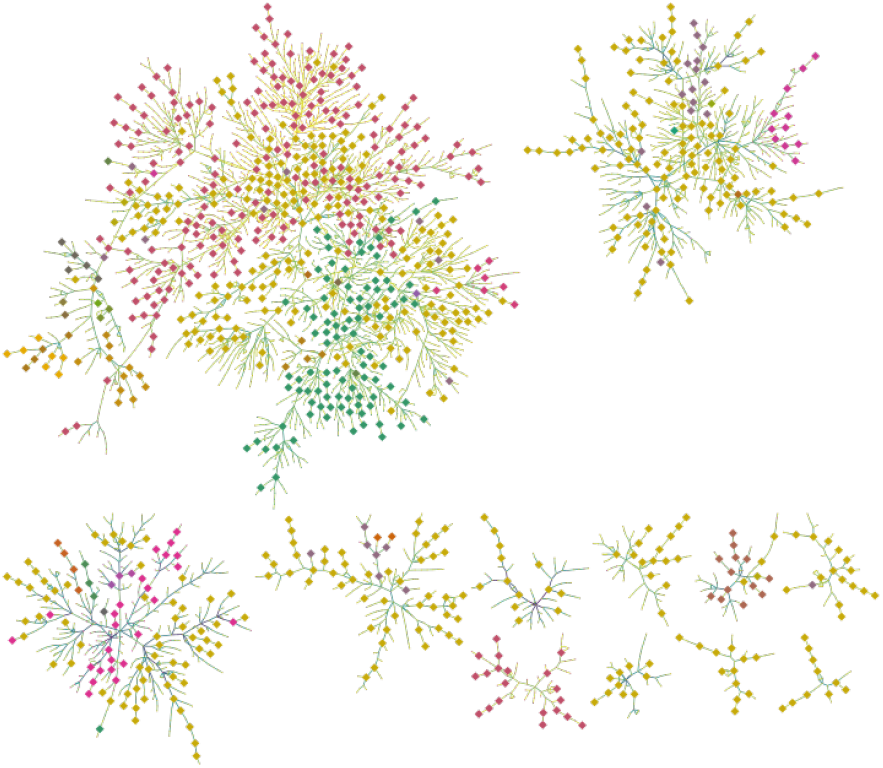
Joint MST network for the 08/01/2021-08/14/2021 time period, filtering on edges with greater than 36% support. Node size indicates origin (large dots are Houston samples), nodes are colored by pango lineage, and edge color indicates patristic distance, where yellow edges are close and blue edges are relatively distant.

**Figure 4:**
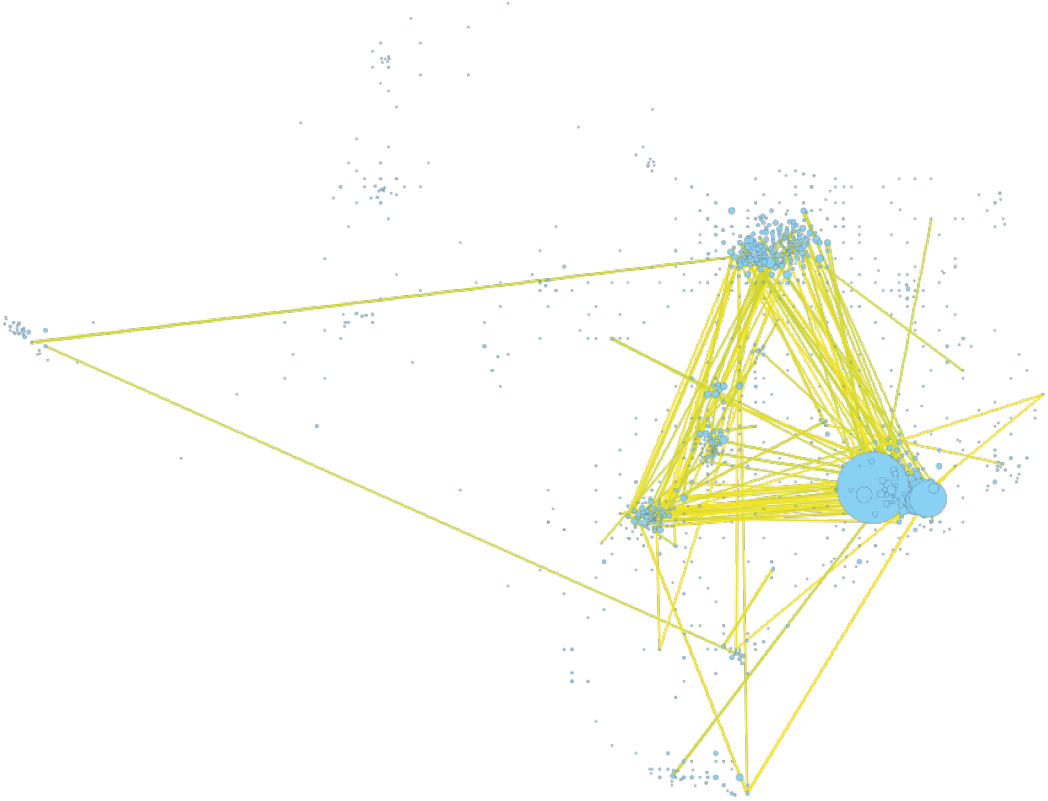
Sample counts by zip code with potential transmission pathways, filtering on patristic distances less than 0.036. Each node represents a zip code and its radius is determined by sample size.

## 4 Conclusions

We created a database for integration of phylogenies and sample metadata which can be used to aid in pandemic response. As we intend for this work to be a proof of concept, there are several limitations we did not address. In addition to the lack of contact tracing and venue affiliation data to corroborate the distance-based transmission estimates, we did not include reference sequences from neighboring regions, preventing us from identifying quantity or origin of variant introductions. While limited in the inferences we can draw from this example use case, future work will integrate contact tracing and venue affiliation data, enabling the development of new methods for transmission network inference.

While this initial analysis could be generated without necessitating a graph database, we benefit at scale with such an approach. Most tools are designed to focus on a single data source or integrate disparate data for individual outbreaks. The ongoing COVID-19 pandemic presents a unique challenge in both variety and volume of data. Graph databases frequently include millions, and in some applications, trillions, of nodes and relationships.[8, 9] Further, many network analysis algorithms, e.g. community detection, similarity, pathfinding, and node embedding, have implementations available in Neo4j. Incorporating additional data layers such as contact tracing and venue affiliation data should enable for more accurate inference of transmission networks and provide opportunities to develop new methods integrating these data. A more immediate next step would be to leverage community detection techniques such as Markov clustering or HDBSCAN to separate cluster signal from noise.[10, 11]

We developed a Python helper script to parse Newick tree files and several Cypher scripts (Neo4j graph database query language) to populate the database, calculate the patristic distances, generate MSTs across the samples, and summarize a joint MST network. https://github.com/glstott/PMeND

## References

[1] Ellsworth M. Campbell, Anthony Alan Boyles, Anupama Shankar, Jay Kim, Sergey Knyazev, Roxana Cintron, and William M. Switzer. Microbetrace: Retooling molecular epidemiology for rapid public health response. PLOS Computational Biology, 2021.

[2] Adriano de Bernardi Schneider, Colby T. Ford, Reilly Hostager, John Williams, John Williams, Michael Cioce, Ümit V. Çatalyürek, Joel O. Wertheim, Daniel Janies, and Daniel Janies. Strainhub: a phylogenetic tool to construct pathogen transmission networks. Bioinformatics, 2019.

[3] Chad Vicknair, Michael Macias, Zhendong Zhao, Xiaofei Nan, Yixin Chen, Yixin Chen, Yixin Chen, Yixin Chen, and Dawn Wilkins. A comparison of a graph database and a relational database: a data provenance perspective. ACM SE’10, 2010.

[4] Stephen A. Smith, Joseph W. Brown, and Cody E. Hinchliff. Analyzing and synthesizing phylogenies using tree alignment graphs. PLOS Computational Biology, 2013.

[5] Ellsworth Campbell, Hongwei Jia, Anupama Shankar, Debra L. Hanson, Wei Luo, Silvina Masciotra, S. Michele Owen, Alexandra M. Oster, Romeo R. Galang, Michael W. Spiller, Michael W. Spiller, Sara J. Blosser, Erika Chapman, Jeremy C. Roseberry, Jessica Gentry, Pamela Pontones, Joan Duwve, Paula Peyrani, Ron M. Kagan, Jeannette M. Whitcomb, Philip J. Peters, Walid Heneine, John T. Brooks, and William M. Switzer. Detailed transmission network analysis of a large opiate-driven outbreak of hiv infection in the united states. The Journal of Infectious Diseases, 2017.

[6] Mathieu Bastian, Sébastien Heymann, and Mathieu Jacomy. Gephi: An open source software for exploring and manipulating networks. ICWSM, 2009.

[7] Paul Shannon, Andrew Markiel, Owen Ozier, Nitin S. Baliga, Jonathan T. Wang, Daniel Ramage, Nada Amin, Benno Schwikowski, and Trey Ideker. Cytoscape: A software environment for integrated models of biomolecular interaction networks. Genome Research, 2003.

[8] Avery Ching, Avery Ching, Sergey Edunov, Maja Kabiljo, Dionysios Logothetis, and Sambavi Muthukrishnan. One trillion edges: graph processing at facebook-scale. Proc. VLDB Endow., 2015.

[9] Christian Theil Have, Lars Juhl Jensen, and Lars Juhl Jensen. Are graph databases ready for bioinformatics. Bioinformatics, 2013.

[10] Stijn van Dongen and Cei Abreu-Goodger. Using mcl to extract clusters from networks. Methods of Molecular Biology, 2012.

[11] Leland McInnes, John Healy, and Steve Astels. Hdbscan: Hierarchical density based clustering. The Journal of Open Source Software, 2017.

